# VeloViz: RNA-velocity informed embeddings for visualizing cellular trajectories

**DOI:** 10.1101/2021.01.28.425293

**Authors:** Lyla Atta, Arpan Sahoo, Jean Fan

## Abstract

Single cell transcriptomic technologies enable genome-wide gene expression measurements in individual cells but can only provide a static snapshot of cell states. RNA velocity analysis can infer cell state changes from single cell transcriptomics data. To interpret these cell state changes as part of underlying cellular trajectories, current approaches rely on visualization with principal components, t-distributed stochastic neighbor embedding, and other 2D embeddings derived from the observed single cell transcriptional states. However, these 2D embeddings can yield different representations of the underlying cellular trajectories, hindering the interpretation of cell state changes. We developed VeloViz to create RNA-velocity-informed 2D and 3D embeddings from single cell transcriptomics data. Using both real and simulated data, we demonstrate that VeloViz embeddings are able to consistently capture underlying cellular trajectories across diverse trajectory topologies, even when intermediate cell states may be missing. By taking into consideration the predicted future transcriptional states from RNA velocity analysis, VeloViz can help visualize a more reliable representation of underlying cellular trajectories. VeloViz is available as an R package on GitHub (https://github.com/JEFworks-Lab/veloviz) with additional tutorials at https://JEF.works/veloviz/.

## 1 Introduction

Current technologies for high-throughput single cell transcriptomics profiling provide a static snapshot of the transcriptional states in individual cells. Still, the continuum of transcriptional states for cells along dynamic processes such as organ development or tumorigenesis can be used to infer how cell states may change over time (Tritschler *et al*., 2019; Saelens *et al*., 2019). Notably, RNA velocity analysis can be applied to infer dynamics of gene expression and predict the future transcriptional state of a cell from single cell RNA-sequencing and imaging data (La Manno *et al*., 2018; Xia *et al*., 2019).

To interpret such cell state changes from RNA velocity analysis, current approaches project the observed current and predicted future transcriptional states onto 2-dimensional (2D) embeddings in order to visualize the putative directed cellular trajectory (La Manno *et al*., 2018; Zywitza *et al*., 2018; Bastidas-Ponce *et al*., 2019; Zhang *et al*., 2019). Previously used 2D embeddings include those derived from principal components (PC), t-distributed Stochastic Neighbor Embeddings (t-SNE), Uniform Manifold Approximation and Projection (UMAP), and diffusion maps (Coifman *et al*., 2005; Maaten and Hinton, 2008; McInnes *et al*., 2018) established using the observed single cell transcriptional states. However, these approaches can yield different representations of the underlying cellular trajectory. Furthermore, in dynamic processes where intermediate cell states are not well represented due to their transient nature or due to technical limitations in sample collection and processing, current 2D embeddings may be unable to capture global relationships between cell subpopulations thereby hindering downstream interpretation of cell state changes (Kester and Oudenaarden, 2018; Weinreb *et al*., 2018). Although alternative non-visual methods such as identifying dynamic driver-genes have been developed to help interpret information from RNA velocity analysis (Bergen *et al*., 2020), visual representation of cellular trajectories remains an important approach to understanding the overall relationships between cell states.

Here, we developed VeloViz to visualize cellular trajectories by incorporating information about each cell’s predicted future transcriptional state inferred from RNA velocity analysis. Using both real and simulated data representing cellular trajectories, we demonstrate that VeloViz embeddings are better able to consistently capture underlying cellular trajectories across diverse trajectory topologies compared to other evaluated methods. Likewise, given simulated cellular trajectories with missing intermediate cell states, we find that VeloViz embeddings are able to more robustly retain the overall cell state relationships in the underlying trajectories compared to other evaluated methods.

## 2 Method

In order to create an RNA-velocity-informed embedding, VeloViz uses each cell’s current observed and predicted future transcriptional states inferred from RNA velocity analysis to represent cells in the population as a graph (Figure 1, Supplementary Information 1).<u>Briefly,</u> starting with spliced and unspliced RNA counts from single-cell RNA-sequencing (scRNA-seq) data or cytoplasmic and nuclear RNA counts from single cell molecular imaging data, the predicted future transcriptional state of cells are inferred using RNA velocity pipelines such as velocyto (La Manno *et al*., 2018) or scVelo (Bergen *et al*., 2020). We then optionally restrict to overdispersed genes (Fan *et al*., 2016) and unit scale each gene’s variance, as well as mean center each gene’s expression for the observed current and predicted future transcriptional states, followed by dimensionality reduction by projecting these observed current and predicted future transcriptional states into a common PC space. Using this reduced dimensional representation of the observed current and predicted future transcriptional states, VeloViz then computes a composite distance 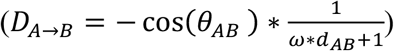 between all cell pairs in the population. The composite distance between two cells, Cell A and Cell B, takes into account: (1) their transcriptional dissimilarity, defined as the Euclidean distance in the common PC space between Cell A’s predicted future state and Cell B’s observed current state (*d*_*AB*_) and (2) their velocity similarity, defined as the cosine correlation between Cell A’s velocity vector and the change vector representing the transition from Cell A to Cell B (*θ*_*AB*_). An additional tuning parameter (*ω*) weighs the relative importance of the transcriptional similarity and the velocity similarity components. In this manner, the composite distance will be minimized when Cell A’s predicted future transcriptional state is similar to Cell B’s observed current transcriptional state and when the direction of Cell A’s RNA velocity is similar to the direction of the transition from Cell A to Cell B. Based on these composite distances, VeloViz creates a k-nearest neighbor graph by assigning *k* directed, weighted edges from each cell to the *k* neighboring cells with smallest composite distances. Edges are further pruned based on parameters that specify the minimum transcriptional and velocity similarity in order to remove spurious cell state relationships. Finally, the pruned graph can be visualized in 2D or 3D using graph layout or graph-embedding approaches such as force-directed layout algorithms (Fruchterman and Reingold, 1991) or UMAP (McInnes *et al*., 2018).

**Figure 1.**
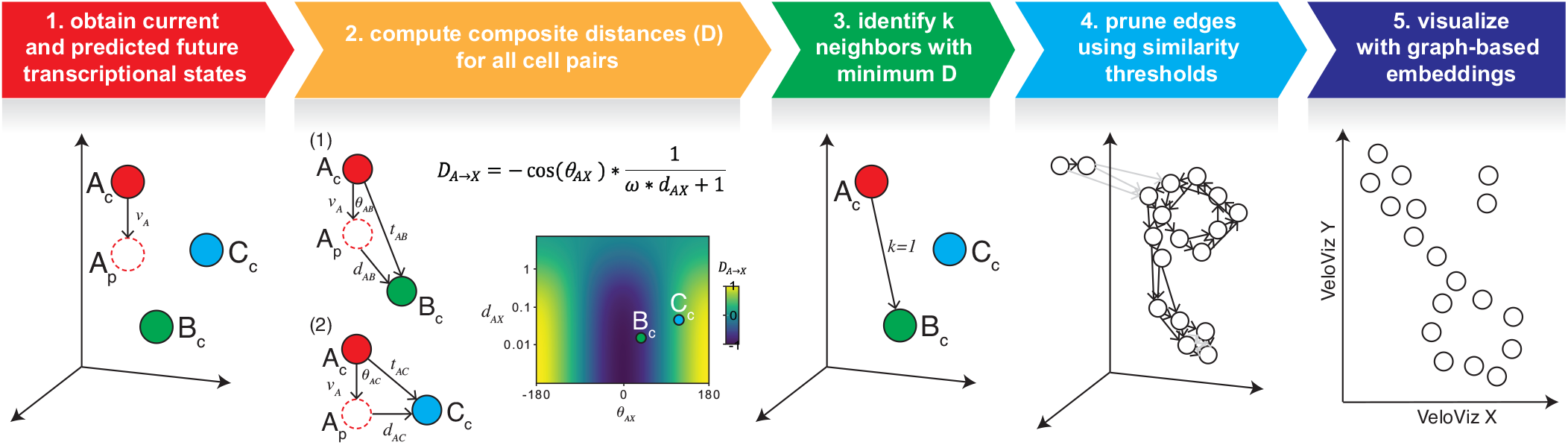
Overview of VeloViz. RNA-velocity informed embeddings are created by VeloViz in five steps: 1) The observed current (X_c_) and predicted future (X_p_) transcriptional cell states are inferred from RNA velocity and reduced into a common PC space; 2) composite distances (D) between all cell pairs are computed. The composite distance from Cell A to Cell X (*D*_*A→X;*_)takes into account the similarity in transcriptional profiles (*d*_*AX*_) between Cell X’s observed current (X_c_) and Cell A’s predicted future transcriptional state (A_p_), and the cosine correlation between Cell A’s RNA-velocity (*v*_*A*_) and the change vector (*t*_*AX*_) representing a transition from Cell A’s current state (A_c_) to Cell X’s current state (X_c_). A distance weight (*ω*) is used to adjust the relative importance of transcriptional similarity and cosine correlation in the composite distance; 3) each cell is represented as a node in a graph, and for each cell, graph edges are assigned to the *k* cells with the minimum composite distances. Edge weights are computed based on composite distances as *weight*_*AB*_ *= max(D) – D*_*AB*_; 4) edges assigned in 3. are pruned (in grey) using transcriptional and velocity similarity thresholds. Edge shade corresponds to edge weight computed based on composite distance, with darker arrows representing edges with larger weights; 5) the resulting graph can be visualized as a 2D or 3D embedding using graph-based embedding approaches.

## 3 Results

### 3.1 Comparing VeloViz to other embeddings

To evaluate the performance of VeloViz, we first assessed VeloViz’s ability to capture cellular trajectories in simulated data representing cycling or branching trajectories (Supplementary Information 2). We compared the VeloViz embeddings to more conventional PC, t-SNE, UMAP, and diffusion map embeddings. To evaluate how accurately each embedding captured the ground truth trajectory, we calculated a trajectory consistency (TC) score (Supplementary Information 3, (Boggust *et al*., 2019)) where high TC scores indicate more accurate representations of the ground truth trajectory. For the simulated cycling trajectory, all evaluated embeddings were able to capture the cycling structure of the trajectory except for the PC embedding (Supplementary Figure 1A). The TC score for the VeloViz embedding was further higher than that of the PC, t-SNE, and UMAP embeddings. For the simulated branching trajectory, the TC score for the VeloViz embedding was higher than TC scores for the t-SNE, UMAP, and diffusion map embeddings (Supplementary Figure 1B-C). Likewise, we evaluated VeloViz’s ability to capture simultaneously cellular trajectories in conjunction with terminally differentiated cell-types using simulated data representing both cycling or branching trajectories with stable a cell population. For the simulated cycling trajectory with a stable cell population, all evaluated embeddings were able to correctly distinguish the cycling and stable populations except for the PC embedding (Supplementary Figure 1 D). Likewise, the VeloViz, t-SNE, UMAP, and diffusion map embeddings preserved the cycling trajectory, while the PC embedding did not. The TC score for the VeloViz embedding was higher than that of the other embeddings. For the simulated branching trajectories with a stable cell population, all embeddings were able to separate the dynamic and stable populations, but only the VeloViz and PC embeddings were able to capture the underlying branching trajectory of the dynamic population (Supplementary Figure 1E-F). This is again reflected in the TC scores, which are consistently higher for the VeloViz and PC embeddings compared to the TC scores for the t-SNE, UMAP, and diffusion map embeddings. These simulation results demonstrate that VeloViz is able to capture trajectories of various topologies compared to other embeddings, which may be better suited for specific topologies.

Next, we assessed VeloViz’s ability to capture cellular trajectories in scRNA-seq data. We applied VeloViz to scRNA-seq data of mouse spermatogenic cells (Supplementary Information 4), where we expect a developmental progression from spermatogonial stem cells to more differentiated spermatids (Hermann *et al*., 2018). For this simple, linear cellular trajectory, VeloViz was able to capture the overall expected trajectory from secondary spermatocytes to early, mid, then late round spermatids (Supplementary Figure 2). Generally, PCA, t-SNE, UMAP, and diffusion map were also able to capture this expected trajectory. To assess VeloViz’s ability to capture more complex trajectory structures, we applied VeloViz to scRNA-seq data of the developing mouse pancreas (Supplementary Information 5), where we expect to see both cycling and branching topologies at different stages of the trajectory. Briefly, we expect cycling ductal cells to give rise to endocrine progenitor-precursor (EP) cells, which become pre-endocrine cells that then differentiate into four hormone producing endocrine cell-types (Alpha, Beta, Delta, and Epsilon cells) (Bastidas-Ponce *et al*., 2019). We observed that while all evaluated embeddings captured the progression of EP cells towards pre-endocrine cells, VeloViz, UMAP, and t-SNE embeddings also captured the terminal branching differentiation into the different endocrine cell-types, which is not clear in the PC or diffusion map embeddings (Figure 2). In addition, VeloViz was better able to capture the cycling structure of ductal cells. Overall, these results indicate that VeloViz embeddings are able to recapitulate expected trends from real scRNA-seq data

**Figure 2.**
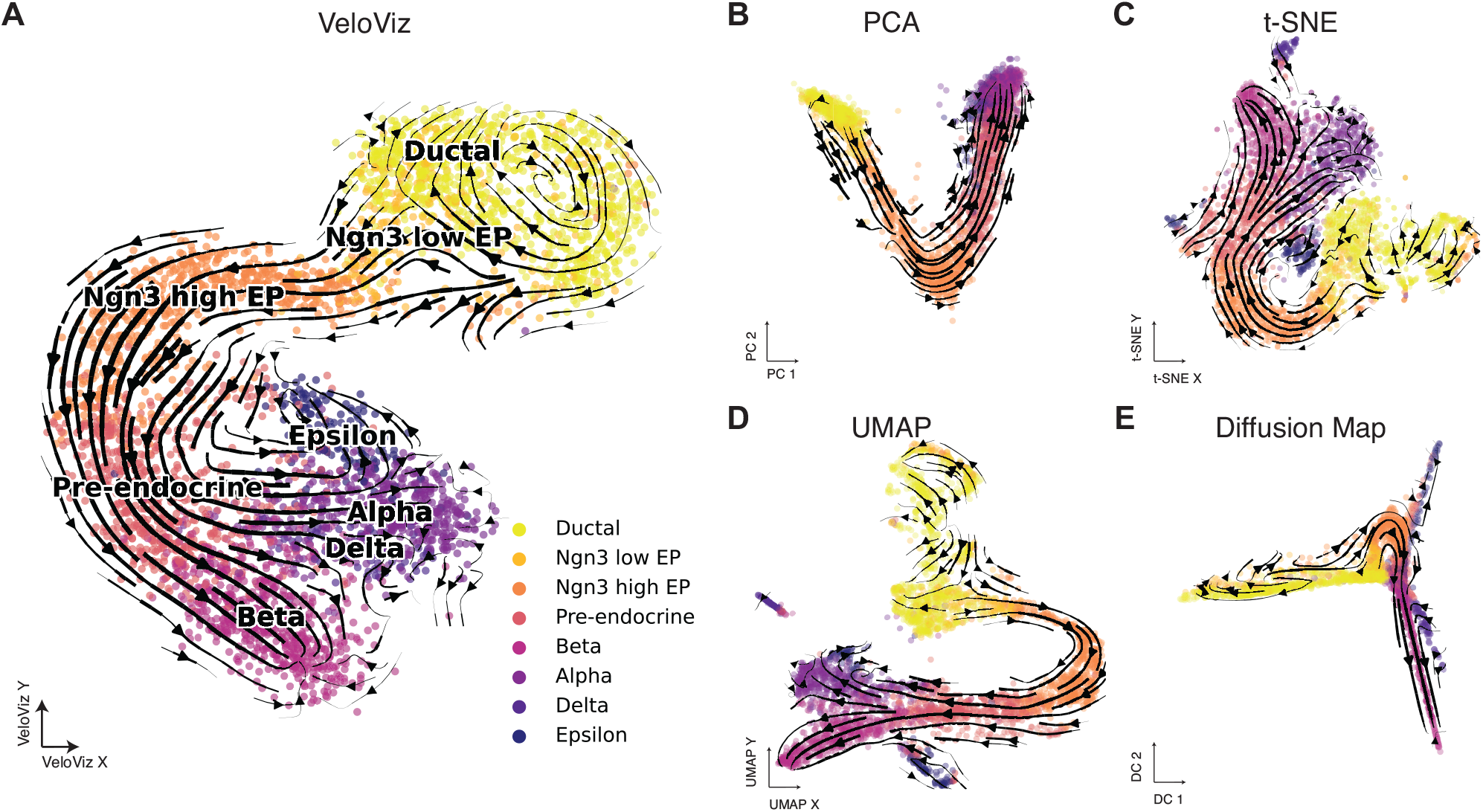
VeloViz reconstructs trajectories from pancreatic endocrinogenesis scRNA-seq. 2D embeddings visualizing pancreatic endocrinogenesis generated using VeloViz (**A**), PCA (**B**), t-SNE (**C**), UMAP (**D**), and diffusion mapping (**E**). Cells are colored by cell state annotations provided in (Bergen *et al*., 2020). Arrows show the projection of velocities derived from dynamical velocity modelling (Bergen *et al*., 2020) onto the embeddings.

To explore the potential of using VeloViz with velocity estimated from other data types, we further applied VeloViz to multiplexed error-robust fluorescent in situ hybridization (MERFISH) data (Xia *et al*., 2019) of cycling cultured U-2 OS cells (Supplementary Information 6). Again, we compared the VeloViz embedding to embeddings constructed using PCA, t-SNE, and UMAP and found that all evaluated embeddings, including VeloViz, were able to capture the expected cycling trajectory (Supplementary Figure 3). In this manner, we find that VeloViz is able to capture cellular trajectories of diverse topologies using both simulated and real data from multiple single cell transcriptomics technologies

### 3.2 Performance with missing intermediate cell states

While uniform sampling of the continuum of transcriptional states for cells along dynamic processes can be used to infer how cell states may change over time, when sampling trajectories with rare or short-lived intermediate cell states or when different cell states are differentially impacted by cell isolation protocols, intermediate cell states may be lost leading to gaps in the observed cellular trajectory (Krishnaswami *et al*., 2016; Villani *et al*., 2017; MacLean *et al*., 2018; Moffitt *et al*., 2018; Slyper *et al*., 2020; Fan *et al*., 2020). We hypothesized that incorporating information about each cell’s predicted future transcriptional state could enable VeloViz to more robustly construct representative cellular trajectories even when the sampled cell states contain missing intermediate cell states or gaps in the underlying trajectory.

To evaluate the robustness of VeloViz in visualizing trajectories with such missing intermediate cell states, we used simulated and real single cell transcriptomics data where some intermediate cells were removed, creating a trajectory gap. Because t-SNE and UMAP preferentially preserve local cell-cell relationships, we hypothesized that these embeddings would result in two distinct clusters of cells before and after the simulated gap (Kobak and Berens, 2019; Heiser and Lau, 2020). Therefore, in addition to TC scores, we calculated a gap distance (Supplementary Information 3), which measures the distance in the 2D embedding space between cells before and after the simulated gap in the trajectory. Embeddings that preserve the underlying trajectory despite this simulated gap will have a smaller gap distance. A small gap distance between cells that are part of the same trajectory will facilitate a clearer depiction of the underlying cell transitions compared to a large gap distance which may erroneously suggest that the cells are unrelated.

Indeed, for the simulated cycling trajectory where cells corresponding to a segment of the cycle were removed (Supplementary Information 2), VeloViz was the only embedding able to clearly represent the cycling structure of the trajectory (Supplementary Figure 4A). The gap distance in the VeloViz embedding was also smaller than in t-SNE, UMAP, and diffusion map embeddings. Likewise, for the simulated branching trajectories where cells corresponding to a segment of an intermediate branch were removed (Supplementary Information 2), only VeloViz and PCA were able to preserve the underlying topology (Supplementary Figure 4B-C). The gap distance in the VeloViz embedding was smaller than that in the t-SNE, UMAP, and diffusion map embeddings. In contrast, t-SNE and UMAP split cells before and after the simulated gap into distinct clusters as expected. TC scores were also consistently higher for VeloViz than with t-SNE, UMAP, and diffusion map embeddings. Similar trends were observed with simulated data that included both dynamic cycling and branching populations with missing intermediate cell states along with a stable cell population (Supplementary Figure 4D-F).

Likewise, for the U-2 OS MERFISH data, to simulate missing intermediate cell states, we removed cells in the G2/M cell cycle phase. Briefly, we identified cells in the G2/M cell cycle phase by computing for each cell a G2/M score based on the aggregated expression of canonical G2/M phase genes (Supplementary Information 6). As before, we compared the VeloViz embedding to those constructed with PCA, t-SNE, and UMAP. We found that VeloViz was better able to retain the cycling trajectory despite the missing G2/M cells compared to the other evaluated embeddings (Supplementary Figure 5).

Similarly, for the developing mouse pancreas scRNA-seq data, to simulate missing intermediate cell states, we removed pre-endocrine cells and used cell latent time (Bergen et al., 2020) to identify cells before and after pre-endocrine cells in the developmental trajectory and to calculate gap distances in the recalculated embeddings (Supplementary Information 5). Notably, while all embeddings depicted the transition from ductal cells to endocrine progenitors, the subsequent transition from endocrine progenitors into terminal endocrine cell-types was best captured by VeloViz. As expected, t-SNE and UMAP split ductal and endocrine progenitor cells from terminal endocrine cell-types, which is reflected in the gap distances (Figure 3). In particular, the position of endocrine progenitors and terminal endocrine cells and the resulting velocity streams may lead to the interpretation that these two cell populations are differentiating in two separate trajectories.

**Figure 3.**
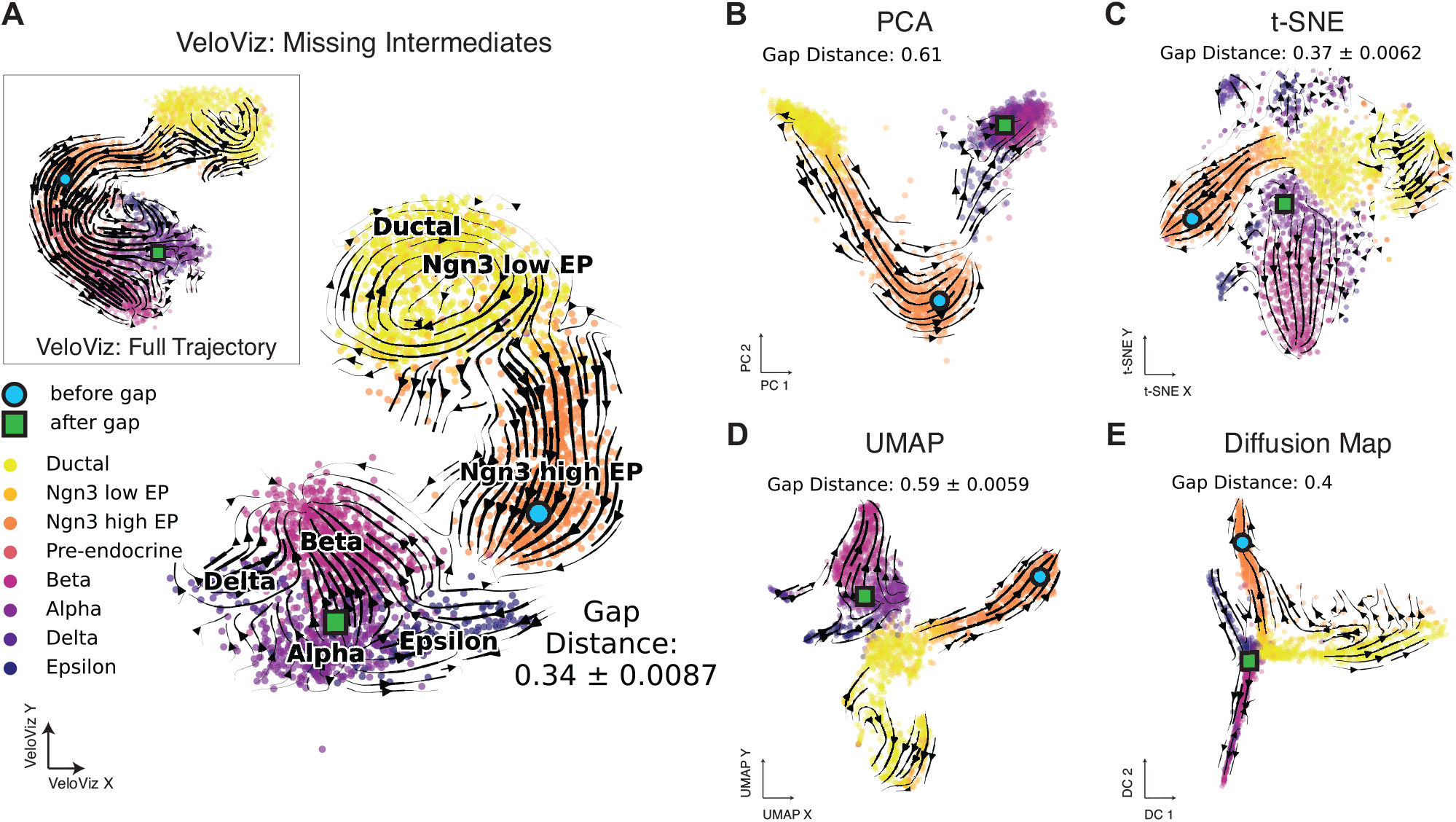
VeloViz reconstructs trajectories from pancreatic endocrinogenesis scRNA-seq data with missing intermediates. **A)** VeloViz 2D embedding visualizing pancreatic endocrinogenesis with pre-endocrine intermediates removed creating a gap in the developmental trajectory. Inset shows the VeloViz embedding of the full dataset. Cells are colored by cell state annotations provided in (Bergen *et al*., 2020). Arrows show the projection of velocities derived from dynamical velocity modelling onto the VeloViz embeddings. Gap distances measure the median distance in the 2D embedding between the 300 cells before and after pre-endocrine cells in the developmental trajectory (Supplementary Information 3iii). Blue circle and green square indicate the median coordinates of cells before and after pre-endocrine cells in the developmental trajectory, respectively. **B-E)** 2D embeddings visualizing pancreatic endocrinogenesis with removed pre-endocrine intermediates using PCA, t-SNE, UMAP, and diffusion mapping, respectively with arrows showing the projection of velocities derived from dynamical velocity modelling.

Still, because low dimensional representations can vary depending on parameter choices, we explored the effect of changing these parameters on t-SNE and UMAP visualizations to see if certain parameter choices would yield visualizations more representative of the underlying cellular trajectory. For t-SNE embeddings, the perplexity parameter affects the extent to which the embedding reflects global vs. local structure, with higher values resulting in embeddings that better preserve global structure (Kobak and Berens, 2019). However, with a gap in the trajectory, the t-SNE embeddings result in two distinct clusters of cells before and after the trajectory gap, even at large perplexity values (Supplementary Figure 3). Likewise, for UMAP, we varied the values of two parameters: minimum distance, which controls how densely packed points are in the embedding with small values resulting in more dense clusters, and the number of neighbors, which functions similarly to perplexity in t-SNE (McInnes *et al*., 2018). As with t-SNE, when embedding data with a simulated gap, UMAP is unable to capture the expected trajectory even at large values of number of neighbors (Supplementary Figure 7). This indicates that when intermediate cell states are missing, t-SNE and UMAP embeddings may be unable to recapitulate the expected underlying trajectory structure regardless of parameter choices. Overall, we find that VeloViz is able to visualize a more reliable representation of underlying trajectories even when intermediate cell states may be missing.

### 3.3 Scalability

Given the increasing availability of large single cell transcriptomics datasets (Lähnemann *et al*., 2020), we sought to evaluate the scalability of VeloViz with increasing cell numbers. Briefly, we down-sampled a dataset of approximately 10,000 cells (10X Genomics, 2020) to create datasets ranging from 100 cells to 9295 cells. For each dataset, we calculated velocity using velocyto.R and constructed an embedding using VeloViz while evaluating runtime and memory usage (Supplementary Information 7). We find that both runtime and memory usage of VeloViz scales linearly with the number of cells and is comparable to that of RNA velocity estimations (Supplementary Figure 8).

## 4 Discussion

In order to facilitate better visual representation of relationships between cell states in single cell transcriptomic data, we developed VeloViz to create low dimensional embeddings that incorporate dynamic information inferred from RNA velocity. We find that VeloViz is able to visualize cellular trajectories of diverse topologies and capture global cell state relationships, even when intermediate cell states may be missing. Particularly when intermediate cell states are missing, we find that visualization with t-SNE and UMAP may result in distinct clusters containing cells before and after trajectory gaps, leading to erroneous interpretation that cells are part of biologically distinct subpopulations rather than the same biological trajectory.

However, several limitations of VeloViz should be considered when using VeloViz embeddings to interpret putative cellular trajectories. Embeddings constructed using VeloViz incorporate multiple user inputted hyperparameters (Supplementary Information 1). We explored the effects of changing these parameters on the visualization of simulated cycling trajectories with missing intermediates and the resulting TC scores (Supplementary Figure 9). We found that the VeloViz embedding was most robust to changes in cosine similarity threshold (t_t_) and most sensitive to changes in *k*. However, without *a priori* knowledge of expected relationships between cell subpopulations, it may be challenging to find the optimal parameter set that yields the most representative embedding. Furthermore, different components of the trajectory being visualized, such as gaps versus branching structures, may have different optimal parameters. Thus, a range of hyperparameters may need to be explored to evaluate the stability of visualized cellular trajectories. Further limitations of VeloViz extend from the limitations of RNA velocity analysis in general. Notably, RNA velocity analysis can only infer cell state changes that are determined by changes in gene expression. Other molecular features such as alternative splicing, chromatin state, post-translational modifications, differential localization, and cell microenvironment that contribute to cell state changes are not considered in RNA velocity analysis, and therefore these cell state changes will not be represented in the VeloViz embedding (Weinreb *et al*., 2018; Tritschler *et al*., 2019). In addition, it remains unknown the degree to which cell state changes are stochastic i.e. the probability that two cells with similar transcriptional states will develop differently. This stochasticity may limit the accuracy of predicting future cell state based on current gene expression dynamics. Ultimately, insights gained from RNA velocity analysis should be considered within the context of other available data, such as differential gene expression, mutational analysis, and targeted experimental validation.

Overall, by taking into account the predicted future transcriptional states of cells from RNA velocity analysis, VeloViz provides an additional approach for visualizing putative cellular trajectories to aid in the interpretation of cellular dynamics from single cell transcriptomics data.

