## Supplementary Information for "VeloViz: RNA-velocity informed embeddings for visualizing cellular trajectories"

#### A. Supplementary Methods

1. VeloViz method
    i. Data preparation
    ii. Composite distance
    iii. Graph construction and layout
    iv. Hyperparameters
2. Simulated data
3. Evaluation metrics
    i. Trajectory consistency score
    ii. Trajectory gap distance
4. Spermatogenesis scRNA-seq data
5. Pancreatic endocrinogenesis scRNA-seq data
6. U-2 OS MERFISH data
7. Scalability

#### B. Supplementary Figures

Supplementary Figure 1: Trajectory reconstruction of simulated data.
Supplementary Figure 2. Directed trajectory reconstruction of spermatogenesis scRNA-seq data.
Supplementary Figure 3. Cell cycle trajectory reconstruction of U-2 OS MERFISH data.

Supplementary Figure 4. Trajectory reconstruction of simulated data with missing intermediate cells.

Supplementary Figure 5. Cell cycle trajectory reconstruction of U-2 OS MERFISH data with missing intermediates.

Supplementary Figure 6. Effects of tSNE perplexity parameter on visualizing pancreas endocrinogenesis.

Supplementary Figure 7. Effects of UMAP minimum distance and number of neighbors parameters on visualizing pancreas endocrinogenesis.

Supplementary Figure 8. Scalability of VeloViz and velocyto.R as a function of cell number.

Supplementary Figure 9. Effects of VeloViz input parameters on visualizing simulated cycling trajectories with missing intermediates.

Supplementary Figure 10. Visualizing VeloViz graphs using alternative layout options.

Supplementary Figure 11. Visualizing VeloViz graphs using UMAP.

**C. Supplementary Tables**

Supplementary Table 1: Parameters used to create 2D embeddings for simulated and real data

**D. Supplementary References**

#### A. Supplementary Methods

1. VeloViz Pipeline

i. Data Preparation

Inputs to VeloViz are the observed current and predicted future transcriptional states inferred from RNA velocity pipelines such as velocyto (La Manno *et al.*, 2018) or scVelo (Bergen *et al.*, 2020). Starting with raw spliced and unspliced scRNA-seq counts, we filter out lowly expressed genes. Alternatively, for MERFISH data, we use cytoplasmic and nuclear RNA counts instead of spliced and unspliced counts, respectively. Then, we calculate RNA velocity using the package velocyto.R to obtain the predicted future transcriptional state. RNA velocity derived from dynamical modeling using the package scVelo can also be used. For dynamical modeling, we obtain the predicted future transcriptional state by adding the velocity output from scVelo to the observed current transcriptional state. We then optionally restrict to overdispersed genes (Fan *et al.*, 2016) and unit scale each gene's variance, as well as mean center each gene's expression for the observed current and predicted future transcriptional states. Finally, we use the observed current transcriptional state matrix to compute PC gene loadings and apply these loadings to both the current and future transcriptional state matrices to project the cells' current and future transcriptional states into a common PC space for downstream calculations.

ii. Composite Distance

VeloViz calculates a composite distance (D) between all cell pairs in the population. This composite distance has two components: (1) a transcriptional similarity component and (2) a velocity similarity component. In the composite distance from Cell A to Cell B, the transcriptional similarity component takes into account the similarity between Cell A's predicted future state and Cell B's observed current state. Transcriptional similarity is defined as the Euclidean distance in PC space ($d_{AB}$) between Cell A's future state and Cell B's current state. The velocity similarity component measures the similarity in transcriptional direction, defined as the cosine correlation, $\cos(\theta_{AB})$, between Cell A's velocity vector and the change vector representing the transition from Cell A to Cell B. Composite distance ranges between -1 and 1. A composite distance of -1 indicates that Cell A's predicted future state is the same as Cell B's observed current state and the direction of Cell A's velocity vector and the change vector from Cell A to Cell B are the same. Conversely, a composite distance of 1 indicates that the direction of Cell A's velocity vector is the opposite of the direction of the change vector from Cell A to Cell B. The relative importance of the transcriptional similarity and the velocity similarity components can be changed via the distance weight parameter ($\omega$). A $\omega$ of 0 results in a composite distance that only takes into account velocity similarity whereas larger $\omega$ places increasing relative importance on transcriptional similarity.

$$D_{A\to B} = -\cos(\theta_{AB}) * \frac{1}{\omega * d_{AB} + 1}$$

##### iii. Graph Construction and Layout

Based on composite distances, VeloViz creates a k-nearest neighbor graph by assigning $k$ directed edges from each cell to the $k$ neighboring cells with smallest composite distances, with edge weights computed based on composite distance as $edge\ weight_{AB} = max(D) - D_{AB}$. Edges are then pruned based on two threshold parameters, a distance threshold ($d_t$) and a similarity threshold ($t_t$). The distance threshold is a quantile threshold for the transcriptional similarity component, computed on all pairwise Euclidean distances $d$ in PC space. Edges between cell pairs where the transcriptional similarity component is greater than the threshold are removed. A distance threshold of 1 includes all edges and does not prune based on transcriptional similarity. The similarity threshold specifies the minimum cosine correlation between the velocity vector and the cell change vector for an edge to be included. A similarity threshold of -1 includes all edges and does not prune based on velocity similarity. The resulting VeloViz graph can be visualized using graph embedding approaches. A 2D embedding using the Fruchterman-Reingold force-directed layout algorithm is visualized by default (Fruchterman and Reingold, 1991). Users also have the option to use the graph output of VeloViz to create higher dimensional 3D embeddings or embeddings using alternative graph layout algorithms (Supplementary Figure 10). Likewise, users can input the nearest neighbor graph returned by VeloViz into alternative graph embedding approaches such as UMAP (McInnes *et al.*, 2018) (Supplementary Figure 11).

##### iv. Parameter options

VeloViz offers multiple user-defined parameter options. These include the number of nearest neighbors for each cell ($k$), distance weighting ($\omega$), percentile distance threshold ($d_t$), and cosine similarity threshold ($t_t$). Based on visualizations of simulated cycling trajectories with missing intermediates across many parameter options (Supplementary Figure 9), we find that the VeloViz embeddings are most robust to changes in cosine similarity threshold ($t_t$) and most sensitive to changes in *k*.

When varying *k*, at small *k* values, VeloViz may produce an embedding where cells before and after trajectory gaps are far apart (Supplementary Figure 9 A-B). This occurs because the Euclidean distances between cells immediately before and after the gap are greater than distances between adjacent cells elsewhere in the cycle. Larger Euclidean distances result in larger composite distances between cells immediately before and after the gap. Therefore, when *k* is small, this results in VeloViz assigning fewer edges from cells before the gap to cells after the gap in favor of assigning edges between adjacent cells on the same side of the gap where composite distance is smaller. In contrast, when *k* is large, VeloViz assigns edges between cells on the same side of the gap as well as between cells on opposite sides of the gap. Because more graph edges between cells results in higher proximity in the 2D embedding, with larger *k*s, cells before and after the gap are closer together (Supplementary Figure 9 E-F).

When varying distance weighting ω, at higher ω, VeloViz may produce an embedding where cells before and after the trajectory gaps are further apart than at lower ω (Supplementary Figure 9 G-L). At higher ω, composite distance is more sensitive to the Euclidean distance between cells. Because cells immediately before and after the trajectory gap are further apart in Euclidean PC space than adjacent cells in other segments of the trajectory, composite distances between them are larger. This leads to (1) fewer edges spanning the trajectory gap and (2) smaller edge weights for edges spanning the trajectory gap. This results in a 2D embedding where cells before and after the gap are further apart.

When varying the distance threshold $d_t$, at lower thresholds, VeloViz may produce an embedding where cells before and after the trajectory gap are further apart (Supplementary Figure 9 M-R). This occurs because the distance-based edge pruning removes edges between cells with large Euclidean distances in PC space. At lower $d_t$ thresholds, when VeloViz removes a larger fraction of graph edges, the remaining edges contain few that span the trajectory gap. This results in a 2D layout where cells before and after the gap are far apart (Supplementary Figure 9 Q-R). In contrast, when the $d_t$ threshold is higher, fewer edges are removed, leaving more edges spanning the trajectory gap, and resulting in a 2D embedding where cells before and after the gap are closer together.

When varying the similarity threshold $t_t$, at higher thresholds, VeloViz may produce an embedding where cells before and after the trajectory gap are further apart. Since the simulated cell velocity vectors represent a small progression in the cycle, the velocity similarity component of the composite distance is greater between cells that are near each other in the cycle. On the other hand, cells immediately before and after the trajectory gap are further apart in the cycle and therefore have lower velocity similarity. Therefore, pruning edges based on velocity similarity removes edges spanning the trajectory gap and results in a 2D embedding where cells before and after the trajectory gap are further apart.

These results show that VeloViz embedding results may vary depending on the choice of parameters, with the optimal choice of parameters likely depending on the features of the data, the underlying structure of process being visualized, as well as the goals of the visualization. For example, optimal choice of *k* will depend on the size of the data: data with fewer cells will require smaller values of *k* to represent relationships between cell populations. To visualize global relationships between cell populations, users should select larger values of *k*, whereas smaller values of *k* will enable visualization of more local relationships. Selecting *k* to be no larger than the size of the smallest cell subpopulation of interest ensures that trajectories in that subpopulation are not subsumed within more global trends across the whole population. Furthermore, if users expect that their data is missing intermediate cell states, optimal parameter values will likely be larger values of *k*, smaller values of ω, and limited or no edge pruning (high $d_t$ and low $t_t$). If the expected trajectory is unknown, selecting small *k*, small ω, and no edge pruning provides a conservative initial approach that prioritizes local structure and limits the impact of potential gaps in the sampled trajectory.

In general, we recommend that VeloViz users explore results with a range of parameter values.

#### 2. Simulation Methods

To assess VeloViz's ability to construct embeddings that recapitulate underlying trajectories, we simulated data by simulating the common PC representation of the current observed and predicted future transcriptional states. To do this based on a defined trajectory structure, we generated 2D trajectories representing cycling and branching topologies and used these as a backbone to generate the simulated common PC representation of the observed current and predicted future gene expression matrices as detailed below.

Cycling Trajectories:

Data was simulated to represent three different trajectory types, a cycle, a branching trajectory with two branch points, and a branching trajectory with three branch points. To simulate the cycling trajectory, we generated a uniformly distributed random variable $t_{ic}$ between 0 and $2\pi$ representing current cells states' position in pseudotime. We then simulated the circular trajectory structure in 2D using the parameters $u_i^1$ and $u_i^2$, as well as simulating an uncorrelated third dimension as follows:

$$u_{ic}^{(1)} = cos(t_i) + U_i^{(1)}$$

$$u_{ic}^{(2)} = sin(t_i) + U_i^{(2)}$$

$$u_{ic}^{(3)} = N_i^{(3)}$$

where $U_i^{(k)}$ are uniformly distributed random variables between $t_i - 0.25$ and $t_i + 0.25$ and $N_i^{(3)}$ is a normally distributed random variable with mean $\mu = 0$ and $\sigma = 1$. Corresponding expression of future cell states was simulated by applying a rotational transformation to the first two dimensions of the observed cells states and adding an uncorrelated third dimension as follows:

$$u_{ip}^{(1)} = cos(\alpha) * u_{ic}^{(1)} - sin(\alpha) * u_{ic}^{(2)}$$

$$u_{ip}^{(2)} = sin(\alpha) * u_{ic}^{(1)} + cos(\alpha) * u_{ic}^{(2)}$$

$$u_{ip}^{(3)} = N_i^{(3)}$$

where $\alpha = 0.1 * \pi$ is the angle by which the cycle was rotated.

To simulate a cycling trajectory with missing intermediates, we ordered simulated observed cells based on their pseudotime and removed the first 100 cells (20% of total cells simulated in full trajectory).

Branching Trajectories:

To simulate the branching trajectories, we adapted methods from (Rizvi *et al.*, 2017). Briefly, we generated a uniformly distributed random variable $t_{ic}$ between 0 and 0.8 representing observed cell states' position in pseudotime. Corresponding pseudotime of future cell states was $t_{ip} = t_{ic} + 0.05$. Then, depending on its pseudotime, we randomly assigned each cell to a branch in the trajectory, defined by a parameters $u_{ic}$ and $u_{ip}$ for current and future branch, respectively. $u_{ic}$ and $u_{ip}$ determined the desired branching structure and were defined as follows:

Three-branch trajectory:

For $t_i < 0.2$, $u_i = 0 + n_i$

For $0.2 < t_i < 0.4$, $u_i = t_i - 0.2 + n_i$ or $u_i = 0.2 - t_i + n_i$

For $t_i > 0.4$, $u_i = t_i - 0.2 + n_i$ or $u_i = 0.2 + n_i$ or $u_i = 0.2 - t_i + n_i$

Four-branch trajectory:

For $t_i < 0.2$, $u_i = 0 + n_i$

For $0.2 < t_i < 0.6$, $u_i = t_i - 0.2 + n_i$ or $u_i = 0.2 - t_i + n_i$

For $t_i > 0.6$, $u_i = t_i - 0.2 + n_i$ or $u_i = 0.2 + n_i$ or $u_i = 0.2 - t_i + n_i$ or $u_i = -0.4 + n_i$

where $n_i$ is a normally distributed random variable with mean $\mu = 0$ and $\sigma = 0.1$. Finally, a third uncorrelated dimension was simulated as a normally distributed random variable with mean $\mu = 0$ and $\sigma = 0.1$.

To simulate trajectories with missing intermediates, we removed a segment of the underlying trajectory. For the two-branch trajectory, we removed the segment between the first and second branch points and for the three-branch trajectory, we removed a portion of the segment between the first branch point and one of the second branch points.

To simulate an additional stable population, we followed the above steps but with $t_{ic} = t_{ip}$ and $u_{ic} = u_{ip}$ generated from a normal distribution with means $\mu = 0.1$ and $\mu = 1$, respectively and $\sigma = 0.1$.

To simulate data with both a stationary population and a dynamic population, we generated current observed and predicted future expression for simulated cycling and branching trajectories as described above. To simulate an additional stable population, we followed the same method but assigned constant pseudotime for all cells in both current observed and future predicted states. With the simulated current observed and predicted future expression matrices, we constructed VeloViz embeddings and compared to PCA, t-SNE, UMAP, and diffusion map embeddings. We computed trajectory consistency scores as described in Supplementary Information 3 to quantitatively compare the embeddings.

3. Evaluation metrics

i. Trajectory consistency score

A trajectory consistency (TC) score was computed to evaluate how accurately an embedding captured the ground truth trajectory. This score was adapted from the local neighborhood similarity score in (Boggust *et al.*, 2019). For each cell we identified the k-nearest neighbors in the ground truth trajectory, $kNN_T$, and in the 2D embedding, $kNN_E$, taking k to be 10% of the population size. We then computed the Jaccard similarity coefficient, $J$, between the two sets of neighbors.

$$J_A = \frac{|\, kNN_T(A) \cap kNN_E(A)|}{|\, kNN_T(A) \cup kNN_E(A)|}$$

The TC score is the median Jaccard similarity of all cells in the embedding. TC scores closer to 1 indicate more accurate representation of the trajectory. Because VeloViz, t-SNE, UMAP, and diffusion map are stochastic, we reevaluated the TC score for each embedding using different seeds. For each embedding, we computed standard errors over 100 runs initiated with different seeds. Standard errors were 0 if the embedding was created using a deterministic method (PC) or if the method always converged to the same lower dimensional embedding.

##### ii. Trajectory gap distance

To evaluate how well an embedding reconstructed a trajectory with missing intermediates, we calculated a trajectory gap distance. The trajectory gap distance measures the distance in the 2D embedding space between the medoid of the cells before the gap and the medoid of the cells after the gap. To enable comparisons across embeddings, we divided the medoid distance by the maximum cell-cell distance in the embedding to generate a normalized gap distance. We note that because the gap distance is computed based on the mediods of cells before and after the trajectory gap, if either or both subpopulations are spread out in the embedding, the gap distance may not reflect qualitative trends illustrated by cell positions and velocity streams.

With real data, where the ground truth trajectory is unknown, we used cell latent time (Bergen *et al.*, 2020) calculated on the full trajectory to identify cells before and after the intermediate cells that were removed. We then computed normalized gap distance as above. As with TC scores, we computed standard errors over 100 runs initiated with different seeds. Standard errors were 0 if the embedding was created using a deterministic method (PC) or if the method always converged to the same lower dimensional embedding.

#### 4. Spermatogenesis scRNA-seq data

Spermatogenesis scRNA-seq data was obtained from (Hermann *et al.*, 2018) via the scRNAseq package (version 2.4.0) (Risso and Cole, 2020). We filtered genes that had fewer than 10 spliced or unspliced counts. Using the filtered spliced and unspliced counts, we calculated RNA velocity using the velocyto.R package with the following parameters: projected time (deltaT) = 1, number of nearest neighbors for smoothing (kCells) = 30, top and bottom quantiles to use for git (fit.quantile) = 0.1, and library scaling factor (mult) = 100. From this, we obtained the observed current expression and predicted future gene expression matrices. We normalized the current and future expression matrices using counts per million normalization and log10 transformed with a pseudocount of 1. We then computed PC loadings on the centered and unit scaled current expression matrix and applied these loadings to both the centered and unit scaled current and future expression matrices to project the cells' current and future transcriptional states to a common PC space with 50 PCs. The current and future transcriptional states in PC space were used to create the VeloViz embedding with parameters detailed in Table 1. The VeloViz embedding was compared to the PC, t-SNE (Rtsne R package, version 0.15), UMAP (uwot R package, version 0.1.8), and diffusion map (destiny R package 3.2.0) embeddings created using the hyperparameters specified in Table 1.

5. Pancreatic endocrinogenesis scRNA-seq data

Pancreatic endocrinogenesis scRNA-seq data was obtained from (Bastidas-Ponce *et al.*, 2019) via the scVelo package (version 0.2.3) (Bergen *et al.*, 2020). We filtered genes that had fewer than 10 spliced or unspliced counts. Using the filtered spliced and unspliced counts, we calculated RNA velocity using the velocyto.R package with the following parameters: projected time (deltaT) = 1, number of nearest neighbors for smoothing (kCells) = 30, top and bottom quantiles to use for git (fit.quantile) = 0.1, and library scaling factor (mult) = 100. From this, we obtained the observed current expression and predicted future gene expression matrices. We normalized the current and future expression matrices using counts per million normalization and log10 transformed with a pseudocount of 1. We then computed PC loadings on the centered and unit scaled current expression matrix and applied these loadings to both the centered and unit scaled current and future expression matrices to project the cells' current and future transcriptional states to a common PC space with 50 PCs. The current and future transcriptional states in PC space were used to create the VeloViz embedding with parameters detailed in Table 1.

We compared the VeloViz embedding to PC, t-SNE, UMAP, and diffusion map embeddings. To create these embeddings, we again normalized gene expression counts to counts per million and log transformed with a pseudocount of 1. We restricted to significantly overdispersed genes, centered and unit scaled the expression matrices, and reduced dimension to 50 principal components. Again, 50 principal components served as the input to create the embeddings using t-SNE, UMAP, and diffusion map with parameters detailed in Table 1. PCs 1 and 2 were used for the PC embedding. Random seed was set to 1 for all stochastic embeddings.

To simulate missing intermediates, we used cell type annotations for cells in the pancreatic endocrinogenesis scRNA-seq data provided in the scVelo package (Bergen *et al.*, 2020) to identify and remove pre-endocrine intermediate cells. We followed the same steps described in 3i. to visualize the remaining cells on VeloViz, PC, t-SNE, UMAP, and diffusion map embeddings.

To explore parameter choices in t-SNE and UMAP visualization, we used the pancreatic endocrinogenesis scRNA-seq data with missing intermediates, down-sampled by a factor of 5 for faster runtime. We constructed embedding using t-SNE (Rtsne R package, version 0.15) and UMAP (uwot R package, version 0.1.8) using varying hyperparameter values. We varied perplexity for t-SNE, and minimum distance and number of neighbors for UMAP across a range of values suggested by the package documentation (Krijthe, 2015; Melville, 2020).

We used the scVelo package to visualize velocity streams on the embeddings in Figures 2 and 3.

6. U-2 OS MERFISH data

Multiplexed error-robust fluorescent in situ hybridization (MERFISH) data of cycling cultured U-2 OS cells was obtained from (Xia *et al.*, 2019). Since this MERFISH experiment did not resolve spliced and unspliced forms of a transcript, we instead used nuclear and cytoplasmic counts to determine RNA velocity as described in (Xia *et al.*, 2019). We constructed the VeloViz embedding as follows. First, we determined cytoplasmic expression to be the difference between the total cell expression and nuclear expression. We then filtered genes with total expression lower than 1. Using nuclear and cytoplasmic expression we calculated RNA velocity using the velocyto.R package with the following parameters: projected time (deltaT) = 1, number of nearest neighbors for smoothing (kCells) = 30, top and bottom quantiles to use for git (fit.quantile) = 0.1, and library scaling factor (mult) = 100. From this, we obtained the observed current expression and predicted future gene expression matrices. We normalized the current and future expression matrices using counts per million normalization and log10 transformed with a pseudocount of 1. We then computed PC loadings on the centered current expression matrix and applied these loadings to both the centered current and future expression matrices to project the cells' current and future transcriptional states to a common PC space with 3 PCs. The current and future transcriptional states in PC space were used to create the VeloViz embedding with parameters detailed in Table 1. We compared the VeloViz embedding to embeddings constructed with PCA, t-SNE, and UMAP. We used the velocyto.R package to visualize velocity streams on the embeddings.

To simulate missing intermediates, we removed 90% of cells in the G2/M phase of the cell cycle based on a G2/M score. To construct the score, we aggregated the expression of canonical G2/M genes for each cell (Whitfield *et al.*, 2002). We randomly removed 90% of cells that had G2/M scores in the top 25% and constructed embeddings with the remainder of the cells as described above.

7. Scalability

To evaluate the runtime and memory scalability of VeloViz with respect to the number of cells in the data, we used scRNA-seq data of mouse embryo neurons with approximately 10 thousand cells (10X Genomics, 2020). We filtered cells with fewer than 100 total spliced or unspliced counts and genes with fewer than 100 total spliced or unspliced counts, retaining 9295 cells and 2926 genes. We then generated three random subsamples for each subsample size ranging from 100 cells to 9295 cells. With each subsample, we calculated velocity using velocyto.R using the parameters projected time (deltaT) = 1, number of nearest neighbors for smoothing (kCells) = 30, top and bottom quantiles to use for git (fit.quantile) = 0.1, and library scaling factor (mult) = 100. We constructed the VeloViz embedding using 50 principal components and all genes with parameters k=15, distance weight=1, similarity threshold=0.1 and distance threshold=0.6. For each subsample, we calculated runtime using the Sys.time function in R and memory usage using the profvis R package (version 0.3.7) (Chang *et al.*, 2020). Evaluations were performed using a 3.2 GHz processor and 32 GB of memory.

### B. Supplementary Figures

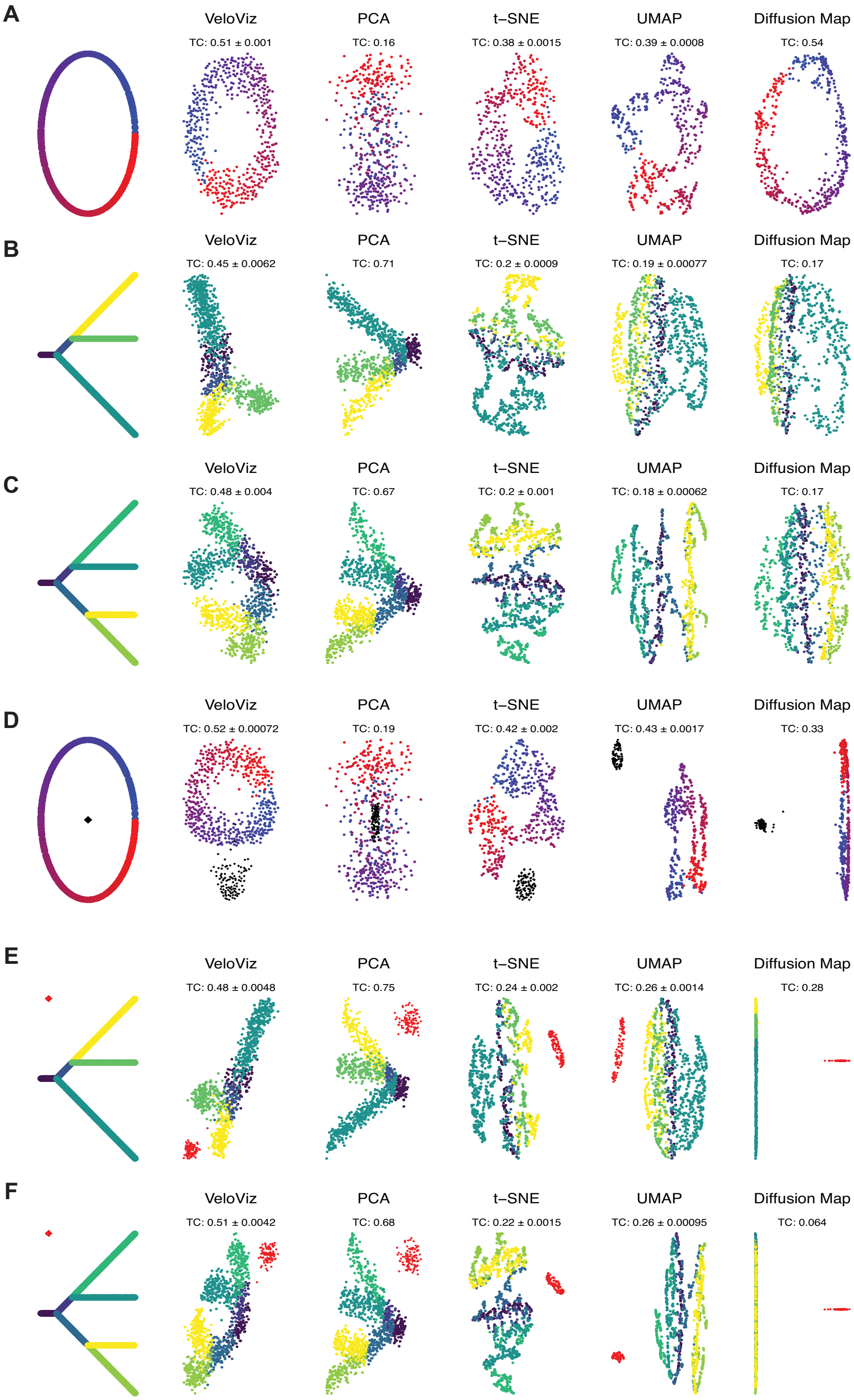

**Supplementary Figure 1: Trajectory reconstruction of simulated data.**

2D embeddings using VeloViz, PCA, t-SNE, UMAP, and diffusion map of simulated **(A)** cycling, **(B)** three-branch, **(C)** four-branch trajectories, and **(D-F)** simulated trajectories with an additional stable population. TC is the mean trajectory consistency score over 100 stochastic runs with different initial seeds.

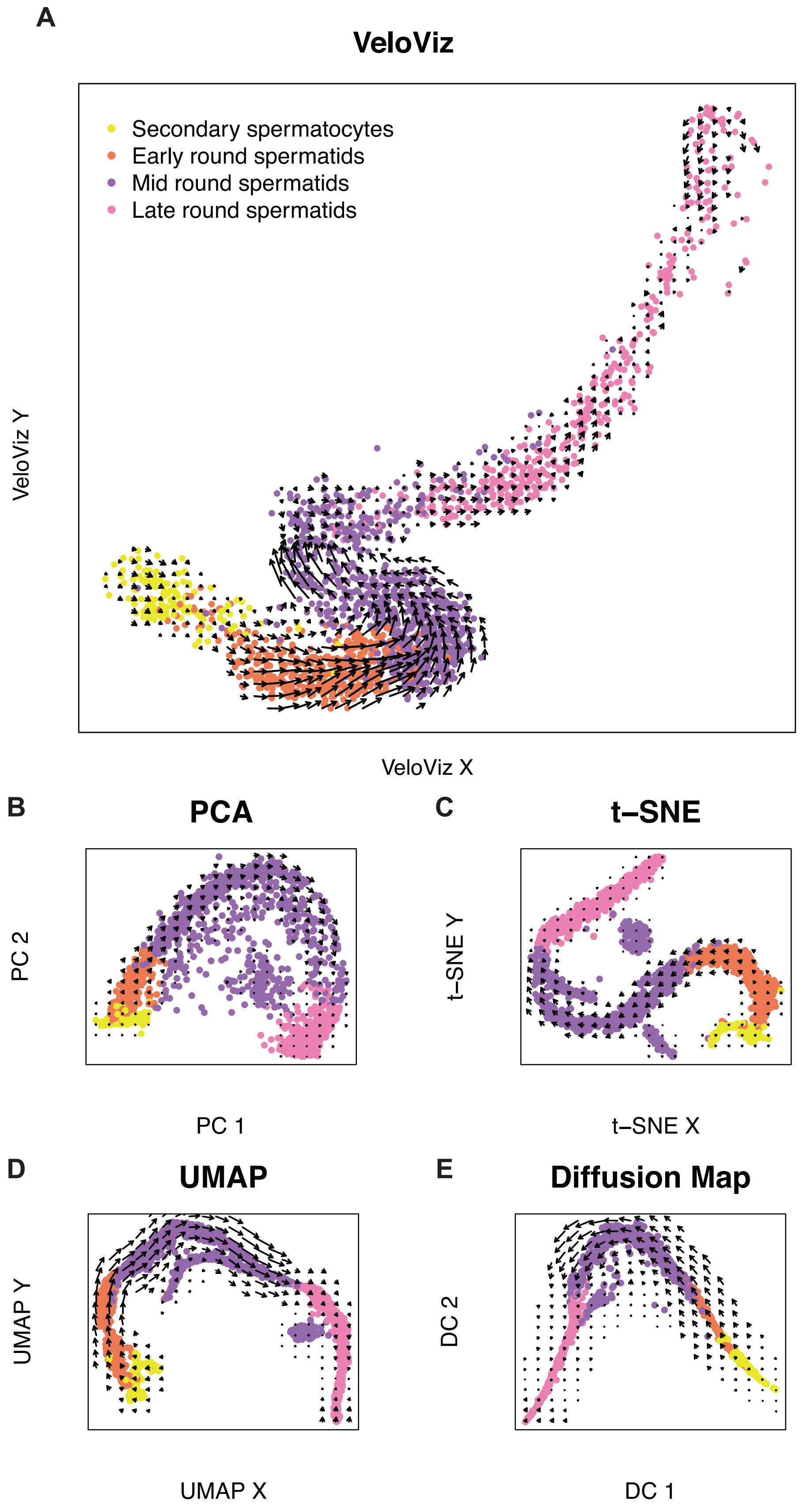
A
VeloViz
Secondary spermatocytes
Early round spermatids
Mid round spermatids
Late round spermatids
VeloViz Y
VeloViz X
B
PCA
PC 2
PC 1
C
t–SNE
t–SNE Y
t–SNE X
D
UMAP
UMAP Y
UMAP X
E
Diffusion Map
DC 2
DC 1

**Supplementary Figure 2. Directed trajectory reconstruction of spermatogenesis scRNA-seq data.** 2D embeddings visualizing spermatogenesis using **(A)** VeloViz, **(B)** PCA, **(C)** t-SNE, **(D)** UMAP, and **(E)** diffusion mapping. Cells are colored by cell type annotations provided in (Risso and Cole, 2020). Arrows show the projection of velocities onto the embeddings by velocyto.R.

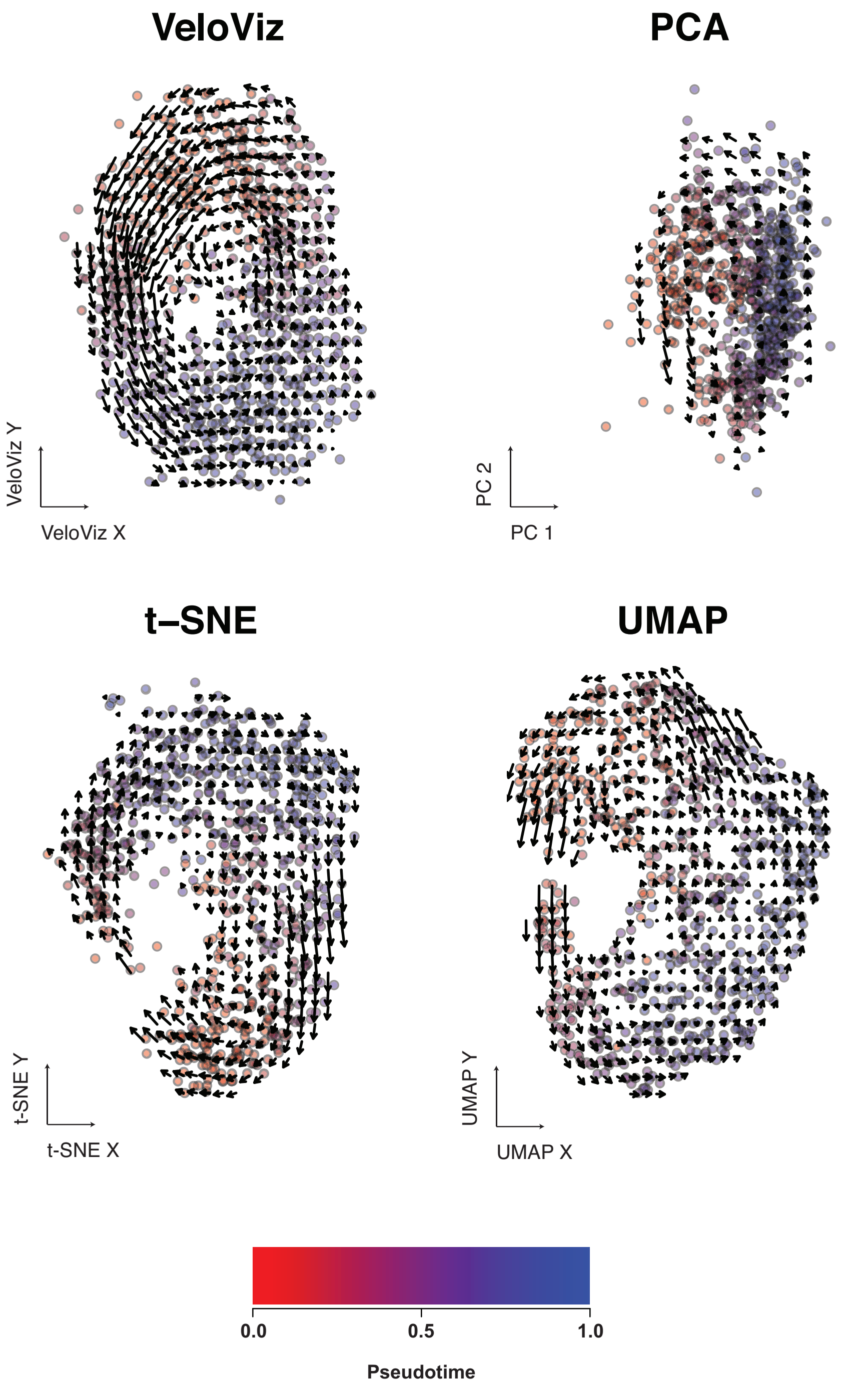


**Supplementary Figure 3. Cell cycle trajectory reconstruction of U-2 OS MERFISH data.** 2D embeddings visualizing cycling cultured U-2 OS cells generated using VeloViz, PCA, t-SNE, and UMAP. Cells are colored by pseudotime. Arrows show the projection of velocities onto the embeddings by velocyto.R.

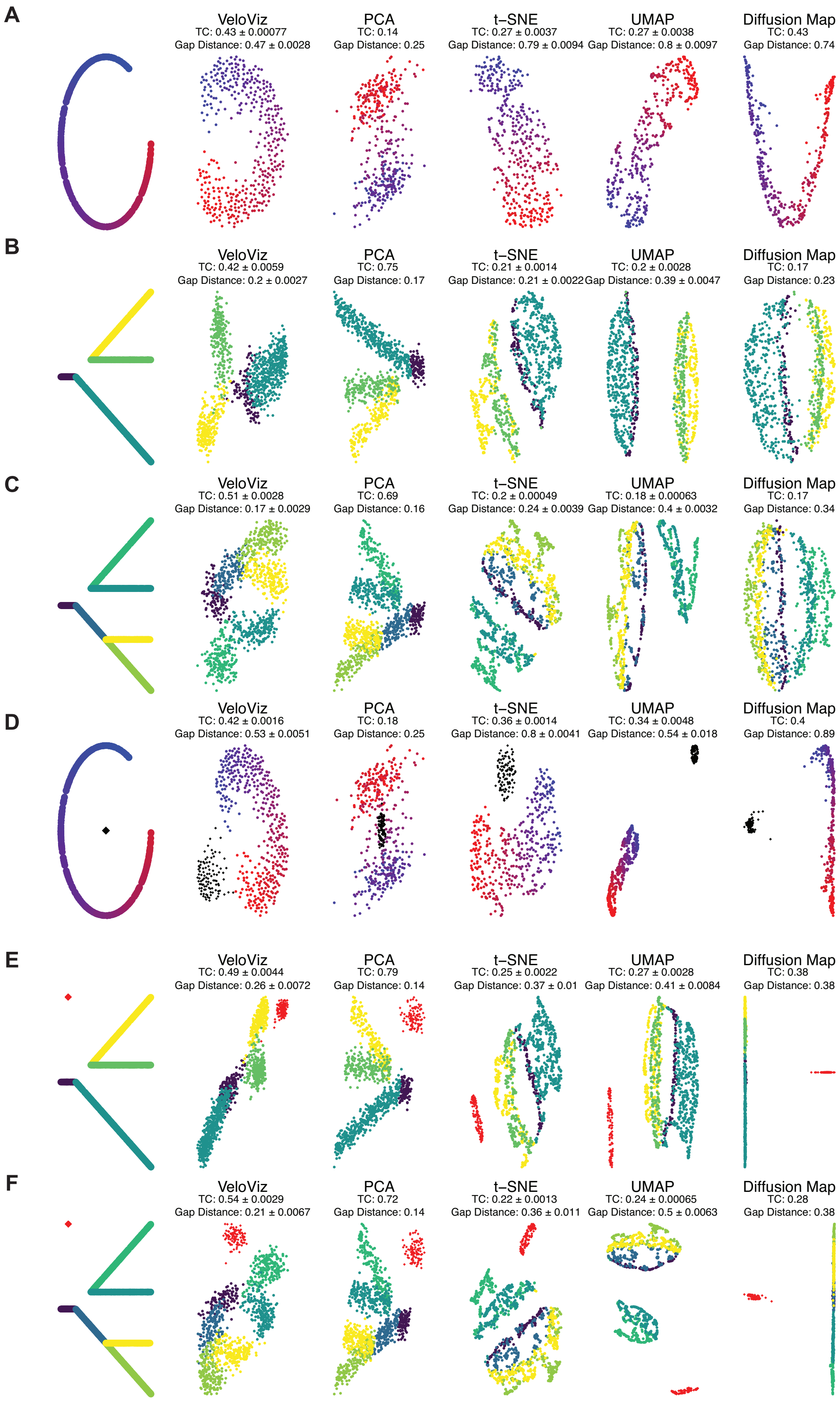
A
VeloViz
TC: 0.43 ± 0.00077
Gap Distance: 0.47 ± 0.0028
PCA
TC: 0.14
Gap Distance: 0.25
t-SNE
TC: 0.27 ± 0.0037
Gap Distance: 0.79 ± 0.0094
UMAP
TC: 0.27 ± 0.0038
Gap Distance: 0.8 ± 0.0097
Diffusion Map
TC: 0.43
Gap Distance: 0.74
B
VeloViz
TC: 0.42 ± 0.0059
Gap Distance: 0.2 ± 0.0027
PCA
TC: 0.75
Gap Distance: 0.17
t-SNE
TC: 0.21 ± 0.0014
Gap Distance: 0.21 ± 0.0022
UMAP
TC: 0.2 ± 0.0028
Gap Distance: 0.39 ± 0.0047
Diffusion Map
TC: 0.17
Gap Distance: 0.23
C
VeloViz
TC: 0.51 ± 0.0028
Gap Distance: 0.17 ± 0.0029
PCA
TC: 0.69
Gap Distance: 0.16
t-SNE
TC: 0.2 ± 0.00049
Gap Distance: 0.24 ± 0.0039
UMAP
TC: 0.18 ± 0.00063
Gap Distance: 0.4 ± 0.0032
Diffusion Map
TC: 0.17
Gap Distance: 0.34
D
VeloViz
TC: 0.42 ± 0.0016
Gap Distance: 0.53 ± 0.0051
PCA
TC: 0.18
Gap Distance: 0.25
t-SNE
TC: 0.36 ± 0.0014
Gap Distance: 0.8 ± 0.0041
UMAP
TC: 0.34 ± 0.0048
Gap Distance: 0.54 ± 0.018
Diffusion Map
TC: 0.4
Gap Distance: 0.89
E
VeloViz
TC: 0.49 ± 0.0044
Gap Distance: 0.26 ± 0.0072
PCA
TC: 0.79
Gap Distance: 0.14
t-SNE
TC: 0.25 ± 0.0022
Gap Distance: 0.37 ± 0.01
UMAP
TC: 0.27 ± 0.0028
Gap Distance: 0.41 ± 0.0084
Diffusion Map
TC: 0.38
Gap Distance: 0.38
F
VeloViz
TC: 0.54 ± 0.0029
Gap Distance: 0.21 ± 0.0067
PCA
TC: 0.72
Gap Distance: 0.14
t-SNE
TC: 0.22 ± 0.0013
Gap Distance: 0.36 ± 0.011
UMAP
TC: 0.24 ± 0.00065
Gap Distance: 0.5 ± 0.0063
Diffusion Map
TC: 0.28
Gap Distance: 0.38

**Supplementary Figure 4. Trajectory reconstruction of simulated data with missing intermediate cells.** 2D embeddings using VeloViz, PCA, t-SNE, UMAP, and diffusion map of simulated cycling **(A)**, three-branch **(B)**, and four-branch **(C)** trajectories with missing intermediates and simulated trajectories with missing intermediates and an additional stable population **(D-F)**. TC: mean trajectory consistency score over 100 stochastic runs with different initial seeds. Gap distances: mean over 100 stochastic runs of the median distance in the 2D embedding between the cells before and after the simulated gap in the trajectory.

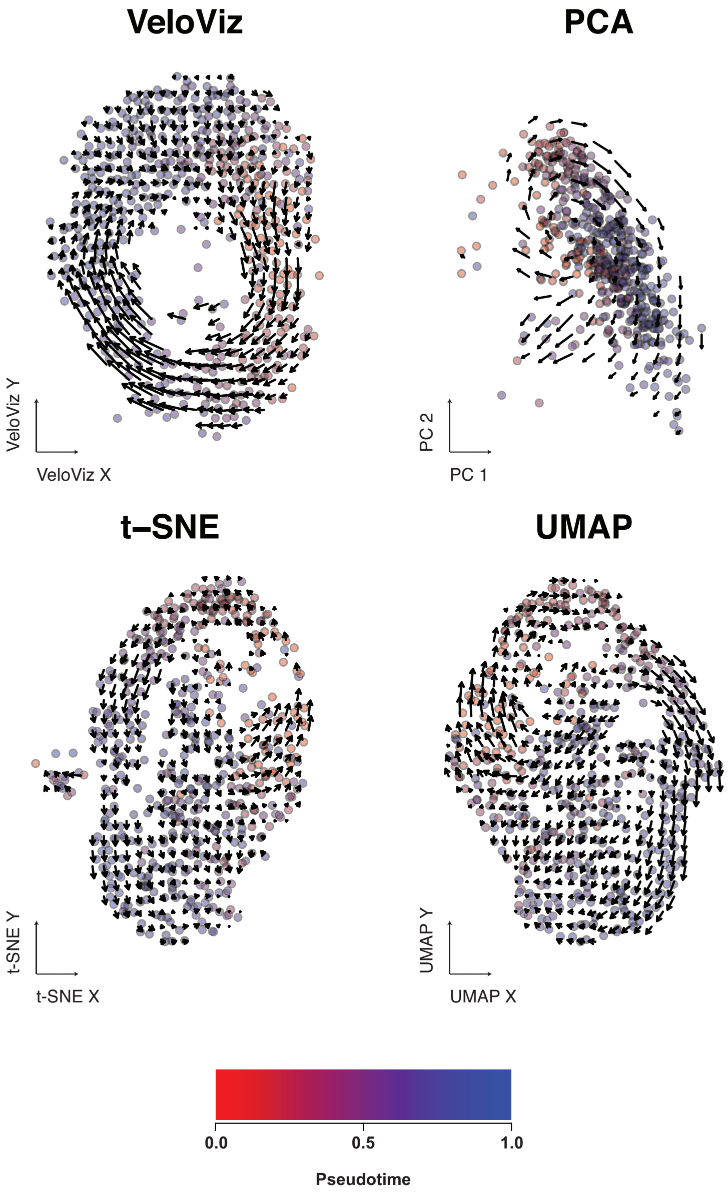


**Supplementary Figure 5. Cell cycle trajectory reconstruction of U-2 OS MERFISH data with missing intermediates.** 2D embeddings visualizing cultured U-2 OS cells with missing intermediates in the G2/M cell cycle phase using VeloViz, PCA, t-SNE, and UMAP. Color indicates pseudotime determined based on the full trajectory. Arrows show the projection of velocities onto the embeddings by velocyto.R.

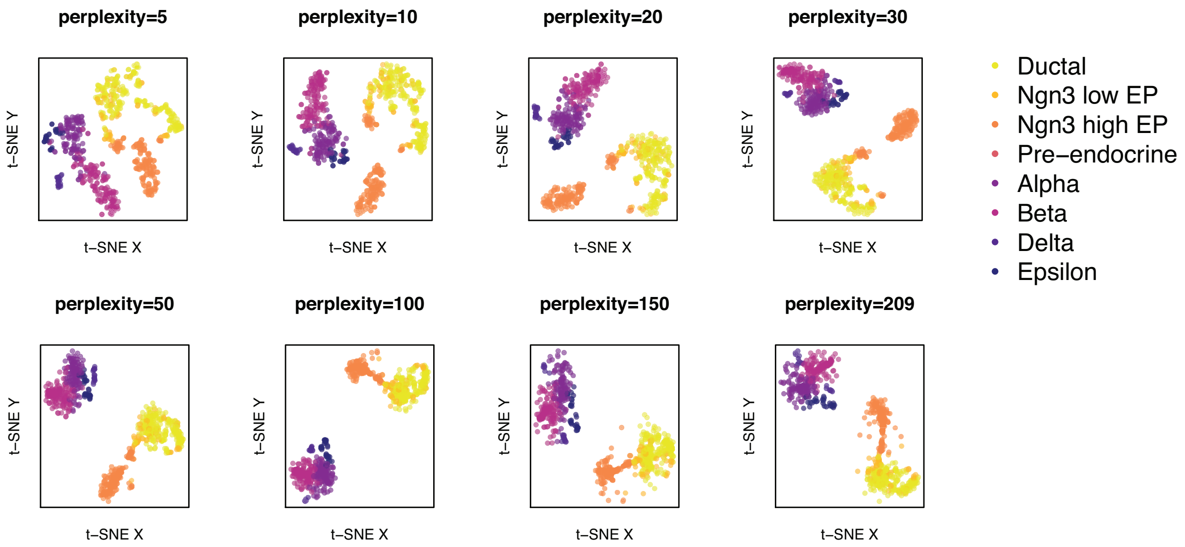


**Supplementary Figure 6. Effects of tSNE perplexity parameter on visualizing pancreas endocrinogenesis.** 2D t-SNE embeddings of pancreatic endocrinogenesis scRNA-seq data with missing intermediates using different perplexity values.

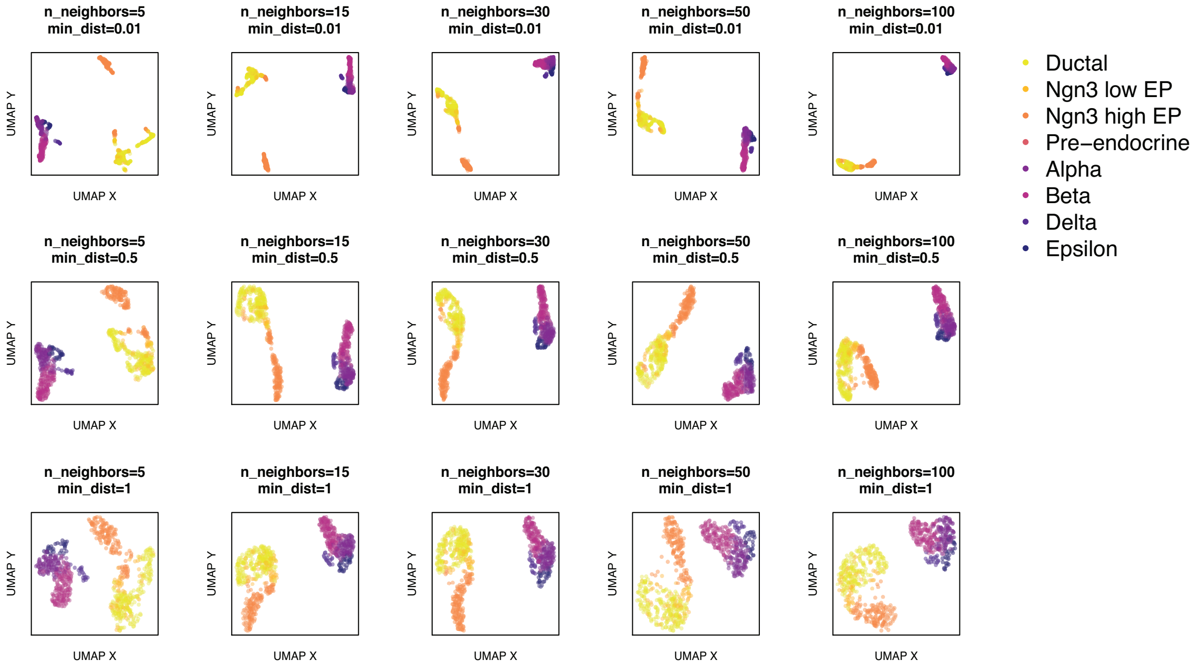


**Supplementary Figure 7. Effects of UMAP minimum distance and number of neighbors parameters on visualizing pancreas endocrinogenesis.** 2D UMAP embeddings of pancreatic endocrinogenesis scRNA-seq data with missing intermediates using different minimum distance and number of neighbors values.

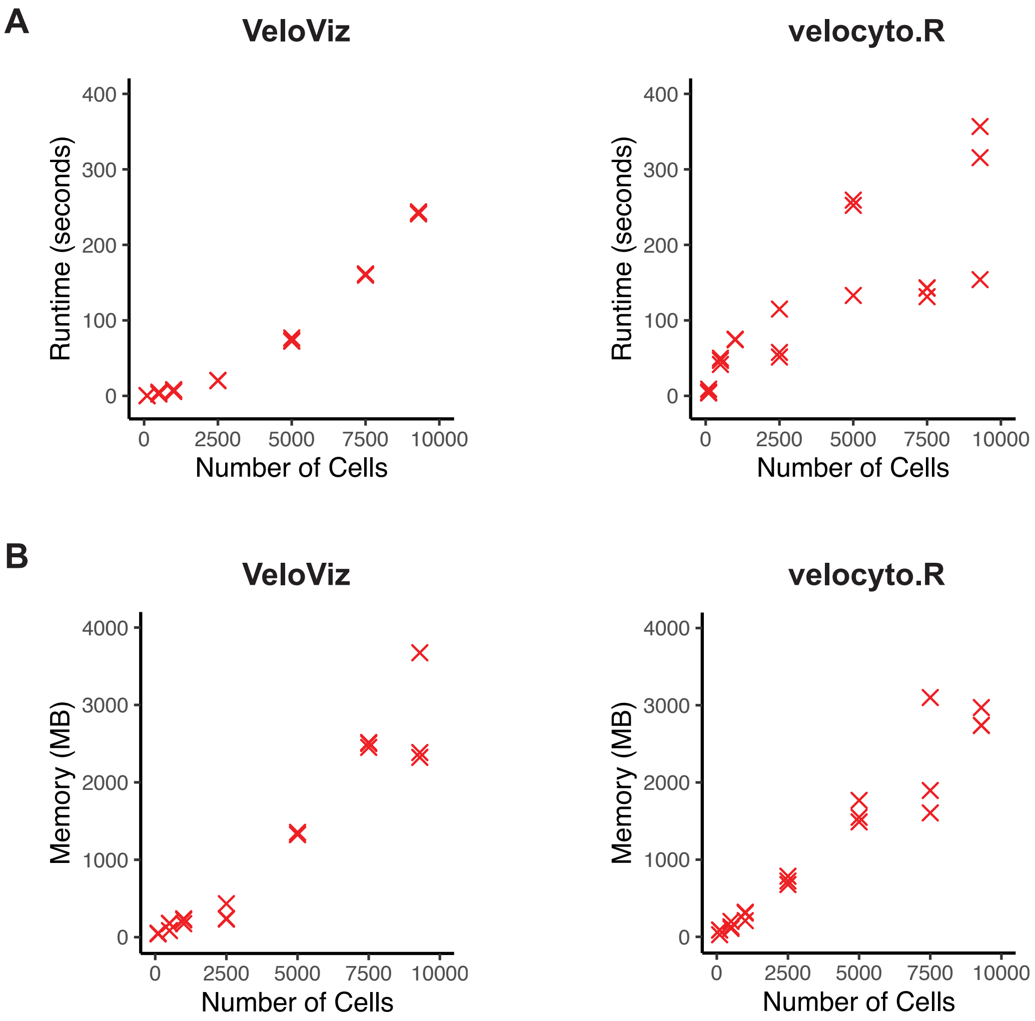


**Supplementary Figure 8. Scalability of VeloViz and velocyto.R as a function of cell number.** Mouse embryo neuron scRNA-seq dataset with approximately 10,000 cells was subsampled creating 3 random samples for a range of subsample sizes (from 100 cells to 10,000 cells). For each subsample size, RNA velocity was computed using velocyto.R and a 2D embedding was constructed using VeloViz. Runtime and memory were evaluated using a 3.2 GHz processor and 32 GB of memory. Scatterplots visualize (A) runtime and (B) memory usage versus the number of cells for both VeloViz and velocyto.R.

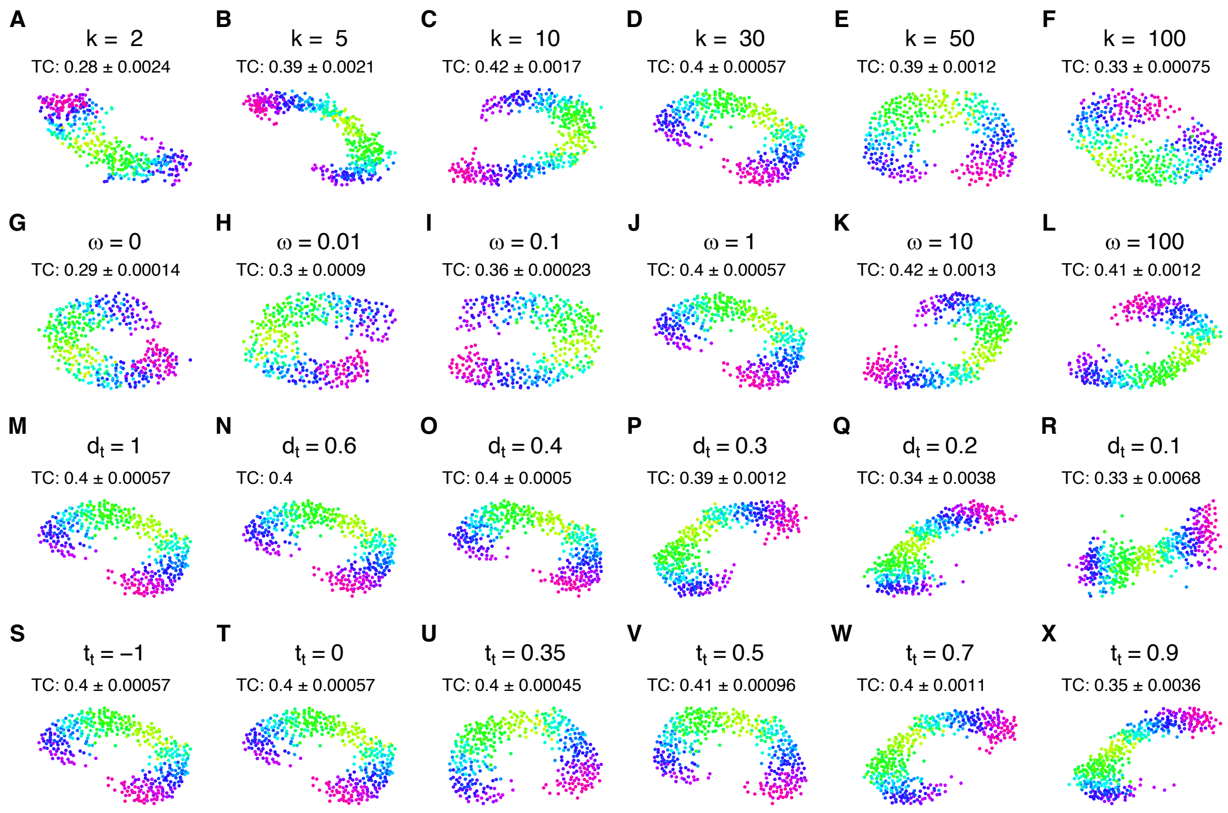


**Supplementary Figure 9. Effect of VeloViz input parameters on visualizing simulated cycling trajectories with missing intermediates. A-F)** VeloViz embeddings with varying number of nearest neighbors for each cell, k, with other parameters constant (ω=1, $d_t$=1, $t_t$=0). **G-L)** VeloViz embeddings with varying weights for the distance component of the composite distance, ω, with other parameters constant (k=30, $d_t$=1, $t_t$=0). **M-R)** VeloViz embeddings with varying percentile distance threshold, $d_t$, with other parameters constant (k=30, ω=1, $t_t$=0). **S-X)** VeloViz embeddings with varying percentile distance threshold, $t_t$, with other parameters constant (k=30, ω=1, $d_t$=1). TC is the mean trajectory consistency score over 100 stochastic runs with different initial seeds.

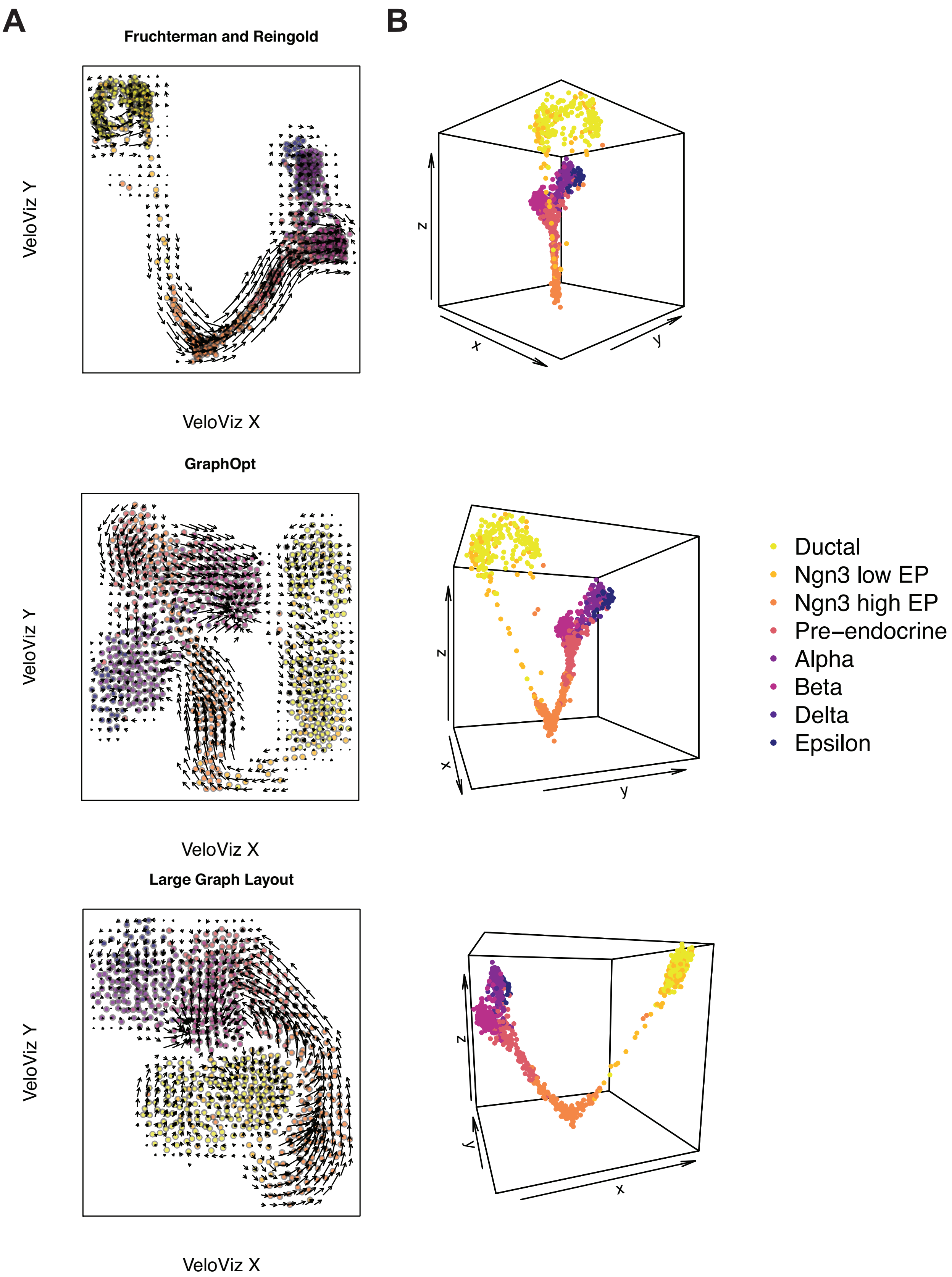
A
B
Fruchterman and Reingold
VeloViz Y
VeloViz X
GraphOpt
VeloViz Y
VeloViz X
Large Graph Layout
VeloViz Y
VeloViz X
z
x
y
Ductal
Ngn3 low EP
Ngn3 high EP
Pre−endocrine
Alpha
Beta
Delta
Epsilon

**Supplementary Figure 10. Visualizing VeloViz graphs using alternative layout options.**

**A)** VeloViz graph of pancreas endocrinogenesis visualized using the Fruchterman-Reingold, GraphOpt, and large graph layout algorithms. Arrows show the projection of velocities onto the embeddings by velocyto.R. **B)** VeloViz graph graph of pancreas endocrinogenesis visualized using Fruchterman and Reingold layout in three dimensions shown from different angles.

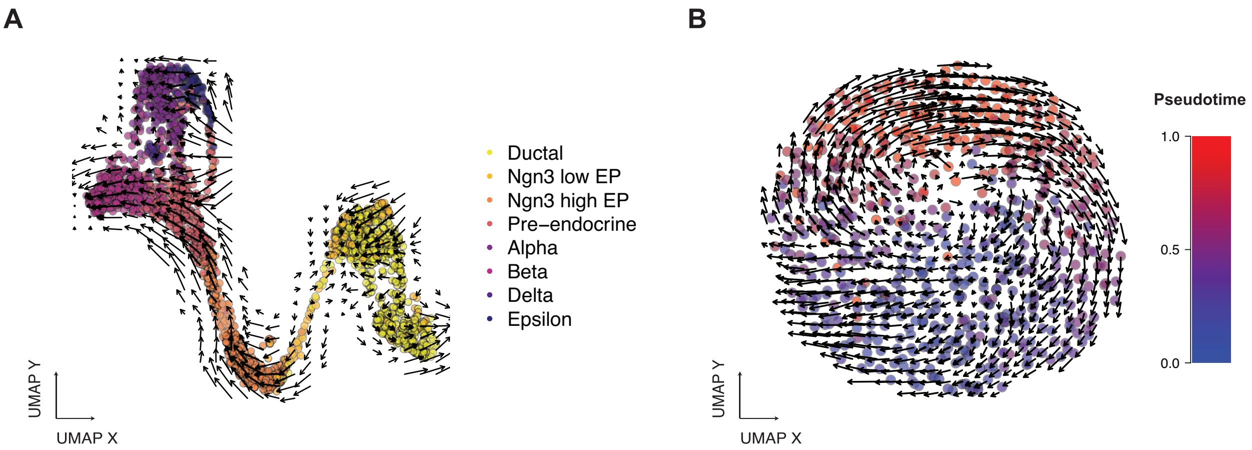


**Supplementary Figure 11. Visualizing VeloViz graphs using UMAP. A)** VeloViz graph of pancreas endocrinogenesis visualized using the UMAP layout. Cells are colored by cell type annotations provided in (Bergen *et al.*, 2020). **B)** VeloViz graph of cycling cultured U-2 OS cells visualized using the UMAP. Color indicates pseudotime. Arrows show the projection of velocities onto the embeddings by velocyto.R.

#### C. Supplementary Tables

**Supplementary Table 1.** Parameters used to create 2D embeddings for simulated and real data including if only overdispersed genes were used (use.ods.genes = True) and the number of PCs (nPCs) used as input.

| | t-SNE<br>perplexity | UMAP | Diffusion Map<br>Number of nearest neighbors | VeloViz |
|---|---|---|---|---|
| Complete cycle<br>nPCs = 3<br>use.ods.genes = False | 100 | n_neighbors = 100<br>min_dist = 0.01 | 50 | $k$ = 30<br>w = 1<br>$d_t$ = 1<br>$t_t$ = 0.25<br>weighted = TRUE |
| Complete three-branch trajectory<br>nPCs = 3<br>use.ods.genes = False | 15 | n_neighbors = 15<br>min_dist = 0.5 | 50 | $k$ = 60<br>w = 10<br>$d_t$ = 0.5<br>$t_t$ = -1<br>weighted = TRUE |
| Complete four-branch trajectory<br>nPCs = 3<br>use.ods.genes = False | 15 | n_neighbors = 15<br>min_dist = 0.1 | 50 | $k$ = 60<br>w = 10<br>$d_t$ = 0.5<br>$t_t$ = -1<br>weighted = TRUE |
| Complete cycle with stable population<br>nPCs = 3<br>use.ods.genes = False | 100 | n_neighbors = 100<br>min_dist = 0.01 | 50 | $k$ = 30<br>w = 1<br>$d_t$ = 1<br>$t_t$ = 0.25<br>weighted = TRUE |
| Complete three-branch trajectory<br>nPCs = 3<br>use.ods.genes = False | 15 | n_neighbors = 15<br>min_dist = 0.5 | 50 | $k$ = 60<br>w = 10<br>$d_t$ = 0.5<br>$t_t$ = -1<br>weighted = TRUE |
| Complete four-branch trajectory<br>nPCs = 3<br>use.ods.genes = False | 15 | n_neighbors = 15<br>min_dist = 0.1 | 50 | $k$ = 60<br>w = 10<br>$d_t$ = 0.5<br>$t_t$ = -1<br>weighted = TRUE |
| Spermatogenesis scRNA-seq<br>nPCs = 50<br>use.ods.genes = False | 30 | n_neighbors = 30<br>min_dist = 0.5 | 100 | $k$ = 5<br>w = 1<br>$d_t$ = 1<br>$t_t$ = 0<br>weighted = TRUE |
| Complete pancreatic endocrinogenesis scRNA-seq<br>nPCs = 50<br>use.ods.genes = True | 30 | n_neighbors = 15<br>min_dist = 0.1 | 100 | $k$ = 25<br>w = 0.05<br>$d_t$ = 1<br>$t_t$ = -1<br>weighted = TRUE |
| Complete U-2 OS MERFISH<br>nPCs = 3<br>use.ods.genes = False | 200 | n_neighbors = 20<br>min_dist = 0.5 | NA | $k$ = 65<br>w = 0.1<br>$d_t$ = 1<br>$t_t$ = -1<br>weighted = TRUE |
| Incomplete cycle<br>nPCs = 3<br>use.ods.genes = False | 100 | n_neighbors = 100<br>min_dist = 0.01 | 50 | $k$ = 30<br>w = 1<br>$d_t$ = 1<br>$t_t$ = 0.25 |

| | | | | |
|---|---|---|---|---|
| | | | | weighted = TRUE |
| Incomplete three-branch trajectory<br>nPCs = 3<br>use.ods.genes = False | 15 | n_neighbors = 15<br>min_dist = 0.5 | 50 | *k* = 60<br>w = 10<br>$d_t$ = 0.5<br>$t_t$ = -1<br>weighted = TRUE |
| Incomplete four-branch trajectory<br>nPCs = 3<br>use.ods.genes = False | 15 | n_neighbors = 15<br>min_dist = 0.1 | 50 | *k* = 60<br>w = 10<br>$d_t$ = 0.5<br>$t_t$ = -1<br>weighted = TRUE |
| Incomplete cycle with stable population<br>nPCs = 3<br>use.ods.genes = False | 100 | n_neighbors = 100<br>min_dist = 0.01 | 50 | *k* = 30<br>w = 1<br>$d_t$ = 1<br>$t_t$ = 0.25<br>weighted = TRUE |
| Incomplete three-branch trajectory with stable population<br>nPCs = 3<br>use.ods.genes = False | 15 | n_neighbors = 15<br>min_dist = 0.5 | 50 | *k* = 70<br>w = 10<br>$d_t$ = 0.5<br>$t_t$ = -1<br>weighted = TRUE |
| Incomplete four-branch trajectory with stable population<br>nPCs = 3<br>use.ods.genes = False | 15 | n_neighbors = 15<br>min_dist = 0.1 | 50 | *k* = 60<br>w = 10<br>$d_t$ = 0.5<br>$t_t$ = -1<br>weighted = TRUE |
| Incomplete pancreatic endocrinogenesis scRNA-seq<br>nPCs = 50<br>use.ods.genes = True | 30 | n_neighbors = 15<br>min_dist = 0.1 | 50 | *k* = 40<br>w = 0.05<br>$d_t$ = 0.95<br>$t_t$ = -1<br>weighted = TRUE |
| Incomplete U-2 OS MERFISH<br>nPCs = 3<br>use.ods.genes = False | 100 | n_neighbors = 20<br>min_dist = 0.5 | NA | *k* = 35<br>w = 0.1<br>$d_t$ = 1<br>$t_t$ = 0.25<br>weighted = TRUE |
